# STRADA deficiency impairs cortical interneuron development in humans and mice

**DOI:** 10.64898/2026.03.30.715326

**Authors:** Ria K. Parikh, Asmaa Hijazi, ThachVu H. Nguyen, Meghna Pandey, Reana Young-Morrison, Danya A. Adams, Sandesh Kamdi, Sophie Tran, Vincent J. Carson, Philip H. Iffland, Louis T. Dang, Peter B. Crino, Whitney E. Parker

## Abstract

Polyhydramnios, Megalencephaly, and Symptomatic Epilepsy syndrome (PMSE/STRADA-related disorder) is a rare neurodevelopmental disorder characterized by megalencephaly (ME), early-onset drug-resistant epilepsy, neurocognitive impairment, and high early mortality, often due to status epilepticus. PMSE is caused by a multi-exon deletion in *STRADA*, encoding STRADA, which regulates the mechanistic target of rapamycin (mTOR) pathway. GABAergic inhibitory interneurons (INs) critically modulate the excitatory:inhibitory balance in cortical and hippocampal networks, and IN deficits contribute to epileptogenesis in several epileptic encephalopathies. However, no studies have investigated INs in PMSE. We used a multimodal approach to study INs in a *Strada*^-/-^ mouse model engineered with the same causative 5-exon deletion identified in human PMSE. We demonstrate that *Strada*/*STRADA* loss causes a reduction of INs in the somatosensory cortex and a corresponding increase in the striatum, representative of remnant ganglionic eminence progenitor origin, in *Strada*-/- mice and a single PMSE brain tissue specimen. RNA sequencing comparing wildtype to *Strada*-/- cortex and striatum corroborated these findings, revealing increased IN-related gene expression (e.g., *Dlx2*) in the striatum and decreased IN-related gene expression (e.g., *Pvalb*) in the developing cortex. Cytoskeletal (e.g., *Tpp3*, *Kank4*, *Map1a*) and mTOR-associated genes (e.g., *Rictor*, *Cryab*) are differentially expressed in the developing cortex, mature striatum, and mature cortex of *Strada*^-/-^ mice. Functional validation confirmed enlarged INs in mouse and human *Strada*/*STRADA*-deficient brain and enhanced S6 phosphorylation in *Strada*^-/-^ striatum. Together, these findings suggest *STRADA*/*Strada* loss contributes to failed IN migration — the first such report in a developmental, mTOR-associated megalencephaly syndrome — highlighting INs as a therapeutic target for seizure prevention in PMSE.

**Key Points:**

- Reduced numbers of cortical inhibitory interneurons were observed in the cerebral cortex of *Strada^-/-^* mice, with striatal interneuron aggregation
- Reduced numbers of cortical inhibitory interneurons, with an aggregation in striatum, were observed in human PMSE brain, supporting the observations in *Strada^-/-^* mouse
- Transcriptomic analysis in *Strada^-/-^* mice reveals evidence of early developmental interneuron and cytoskeletal dysfunction
- We introduce a loss of cortical interneurons as a salient feature of PMSE developmental pathogenesis, potentially contributing to a loss of inhibitory modulation
- This is the first study proposing interneuron migration impairment in the developmental pathogenesis of an mTOR-associated megalencephaly syndrome

## INTRODUCTION

Polyhydramnios, Megalencephaly, and Symptomatic Epilepsy syndrome (PMSE/STRADA-related disorder) is characterized by epilepsy, neurocognitive impairment, and high perinatal and early childhood mortality. PMSE is caused most commonly by a homozygous deletion of exons 9-13 of the *LYK5/STRADA* gene on chromosome 17, resulting in a loss of function of the STE20-related kinase adaptor protein alpha (STRADA).^1^ STRADA is a pseudokinase that activates liver kinase B1 (LKB1) and in turn, adenosine monophosphate-activated protein kinase (AMPK)-driven inhibition of mechanistic target of rapamycin (mTOR) signaling, acting upstream of TSC1/TSC2/TBC1D7 in response to cellular energy depletion.^2^ Loss of STRADA results in mTOR hyperactivity, evidenced by increased phosphorylation of S6 protein (P-S6) in both *Strada* knockout (KO; *Strada^-/-^*) mouse and PMSE human cortex.^3^ mTOR hyperactivation has been repeatedly demonstrated in tuberous sclerosis complex (TSC), focal cortical dysplasia and hemimegalencephaly (mTORopathies).^4–6^ Defining the impact of STRADA loss and mTOR dysregulation on early brain development is therefore critical to understanding PMSE pathogenesis and informing targeted therapies for this and other mTORopathies.

We previously demonstrated that Strada loss impairs migration of cortical excitatory neural progenitor cells (NPCs) in mice.^3,7^ Histopathological analysis of *Strada^-/-^*mice revealed impaired lamination of late-born (Cux1+) excitatory neurons, with aberrant accumulation of these layer 2/3-fated neurons in deeper cortical layers at postnatal day 9 (P9).^8^ *Strada* knockdown *in vitro* in P0 excitatory NPCs and PMSE patient fibroblasts demonstrated mTOR-dependent impairment in cell motility with altered cytoskeletal dynamics, suggesting that altered cortical lamination in PMSE arises from migration defects driven by mTOR hyperactivity.^7^ These findings motivated a pilot clinical trial of the mTOR inhibitor sirolimus (rapamycin) in pediatric PMSE patients, which initially showed promise as a novel and effective epilepsy treatment in this population.^7^ However, seizure frequency returned to baseline over time, motivating a search for additional contributory disease mechanisms.

GABAergic inhibitory interneurons (INs) comprise approximately 20-25% of cortical neurons and are essential for maintaining the excitatory:inhibitory network balance through modulation of excitatory neuron activity.^9,10^ Disrupted IN development, particularly migration, has been shown in multiple developmental epileptic encephalopathies, however, the role of STRADA in IN development and its contribution to PMSE pathogenesis remain unknown^10^.

During brain development, approximately 90% of inhibitory NPCs arise in the ganglionic eminence (GE) before migrating tangentially to the cortex.^11,12^ The GE is divided into medial (MGE) and caudal (CGE) subregions. Parvalbumin (PV)- and somatostatin (SST)-positive INs, the two most abundant cortical IN classes, originate in the MGE, whereas serotonin receptor 3a (5HT3aR)-positive INs originate in the CGE.^13^ While MGE-derived NPC migration is better-characterized than CGE-derived NPCs, recent studies suggest shared migratory features.^14^ In human, tangential migration of inhibitory NPCs begins in the second trimester and continues postnatally.^15,16^ Given that Strada loss disrupts cytoskeletal organization, a process fundamental to cell migration, we hypothesized that STRADA likely plays a key role in IN migration.

Here, we present the first histopathological analysis of INs in PMSE patient and *Strada*^-/-^ mouse brain. We demonstrate that STRADA loss is associated with evidence of impaired migration of inhibitory progenitors, paralleling known defects in excitatory NPC migration.^7,8^ Using multimodal histological approaches, we examine IN subtype expression in human PMSE and *Strada^-/-^* mouse cortex and subcortical remnant GE progenitor regions. We also profile regional mRNA expression in cortex, hippocampus, and subcortical structures of developing (P3) and mature (P21) *Strada* KO and WT mice, focusing on pathways regulating IN development, migration, cytoskeletal organization, and mTOR signaling. Our findings indicate that in both human and mouse, STRADA/Strada loss is associated with reduced cortical IN numbers and retention of INs in subcortical remnant GE regions, consistent with disrupted migration, particularly affecting MGE-derived subtypes. Together, these results identify STRADA as a critical regulator of IN migration and lamination during corticogenesis, highlighting early developmental processes that must be considered in future PMSE disease-modifying therapeutic strategies.

## MATERIALS AND METHODS

### Animals

WT (*Strada^+/+^*), heterozygous (*Strada^+/-^*), and KO (*Strada^-/-^*) mice of both sexes were assayed. The *Strada ^-/-^*mouse line was previously generated in the Crino lab using a neomycin cassette to model the homozygous deletion of *STRADA* exons 9-13 identified in the Old Order Mennonite PMSE founder mutation, replicated in the murine *Strada* ortholog.^8^ As previously reported, *Strada^-/-^* mice exhibit megalencephaly and high pre- and perinatal lethality rates.^8,17^ For histopathology, P21 mice (3-5 per genotype) were analyzed. For bulk RNA sequencing, WT and KO P3 and P21 mice were used (3 per genotype, per age). All procedures were approved by the University of Maryland, Baltimore Institutional Animal Care and Use Committee (AUP-00000032).

### Histopathology and Immunohistochemistry (IHC)

Human pediatric control cortex, hippocampus, and subcortical tissue were obtained from the NIH tissue repository and processed in parallel with PMSE tissue (see “Results”) as previously described.^1,3^ Sections were immunolabeled using DAB-based IHC for GABAergic IN markers. Detailed protocols are provided in **Supplemental Methods.**

At P21, mice were perfused with PBS and 4% paraformaldehyde. Brains were paraffin-embedded and sectioned coronally at 10 µm. Coronal sections underwent standard immunofluorescent labeling with primary antibodies against IN markers, followed by fluorophore-conjugated secondary antibodies and DAPI nuclear counterstaining. Antibody details are provided in **Supplemental Table 1** and **Supplemental Methods.**

### Imaging, Quantification, and Statistics

Human tissue was imaged under consistent brightfield settings. Semi-quantitative analysis and automated particle counting for cell counts were performed in ImageJ following thresholding. Group comparisons for cell size were made using a Kolmogrov-Smirnov test of normality and a Mann-Whitney U test for data sets with a small sample size and non-normal distribution.

Mouse sections were imaged using a Keyence BZ-X710 fluorescence microscope and human sections were imaged using a Keyence BZ-X800 microscope. For each marker, three sections per animal were analyzed, with regions of interest defined using the Allen Brain Atlas. Cells were manually counted in ImageJ based on morphology and nuclear colocalization.^18^ Counts were averaged per animal and analyzed using one-way ANOVA with Tukey’s post-hoc correction. Statistical significance was defined as p<0.05. For the laminar analysis, the cortex was divided into upper (layer 2-3), middle (layer 4), and lower (layer 5-6) thirds and each section was counted and compared separately across genotypes using a one-way ANOVA with Tukey’s multiple comparisons. Cell size in the striatum was measured by tracing the soma of 10 representative cells per section and comparing the average soma size per animal across genotypes using a one-way ANOVA with Tukey’s multiple comparisons.

### Bulk RNA Sequencing (RNAseq)

Cortex, hippocampus, and subcortical tissue (including basal ganglia and thalamus, representative of remnant GE) were collected from WT and KO mice at P21; cortex alone was collected at P3. Tissue was processed for bulk RNAseq at our Institute for Genome Sciences Core. Differential expression analysis was performed using DESeq2, stratified by brain region and genotype. A prospective curated set of 193 genes related to interneuron development, migration, cytoskeletal organization, and mTOR signaling was analyzed by functional category. Full sequencing and bioinformatics methods are provided in **Supplemental Methods.**

## RESULTS

We performed a multimodal histopathological analysis of IN subtype expression in PMSE human brain tissue and *Strada*^-/-^ mouse across cortex, hippocampus, and striatum (serving as the remnant GE progenitor region), representing destination and origin regions for tangential IN migration during neurodevelopment. In parallel, we conducted unbiased transcriptomic analyses of WT and *Strada*^-/-^ mouse cortex at P3 and P21, as well as P21 hippocampal and subcortical (basal ganglia and thalamus, remnant GE) regions. Concordant histopathological and RNAseq findings and cytomegaly across species support the validity the *Strada*^-/-^ mouse as a translationally relevant model of PMSE and support the hypothesis that STRADA loss disrupts IN migration.

### PMSE brain exhibits reduced cortical and hippocampal INs, with increased retention in the striatum

Histopathology from a post-mortem PMSE patient brain (female, 7 months, reported previously in Puffenberger et al., 2007) was compared with age-matched pediatric control tissue. We quantified total INs (GABA+, GAD65+), major IN subtypes (PV, SST, 5HT3aR), and MGE-derived progenitors (NKX2.1) in frontal cortex, hippocampus, and striatum (**Supplemental Figure 5**). These subtypes were selected because they comprise the majority of cortical and hippocampal INs and are strongly associated with epilepsy.^13,19,20^ Semi-quantitative analysis was performed using automated ImageJ-based counting following consistent thresholding (**Supplemental Table 2**).

In PMSE cortex, GABA+, GAD65+, GAD1/67+, PV+, and 5HT3aR+ neurons were reduced relative to control tissue, confirmed by semi-quantitative n=1 analysis **(Figure 1, Supplemental Figure 1).** In the hippocampus, GABA+, 5HT3aR+, and SST+ neurons were decreased. In contrast, the striatum exhibited *increased* numbers of total INs and all major subtypes. NKX2.1+ cell numbers did not differ across brain regions, indicating that reduced cortical and hippocampal INs were not attributable to impaired progenitor differentiation. Together, cortical depletion with striatal accumulation, potentially reflecting remnant GE, supports disrupted IN migration in PMSE.

**Figure 1:**
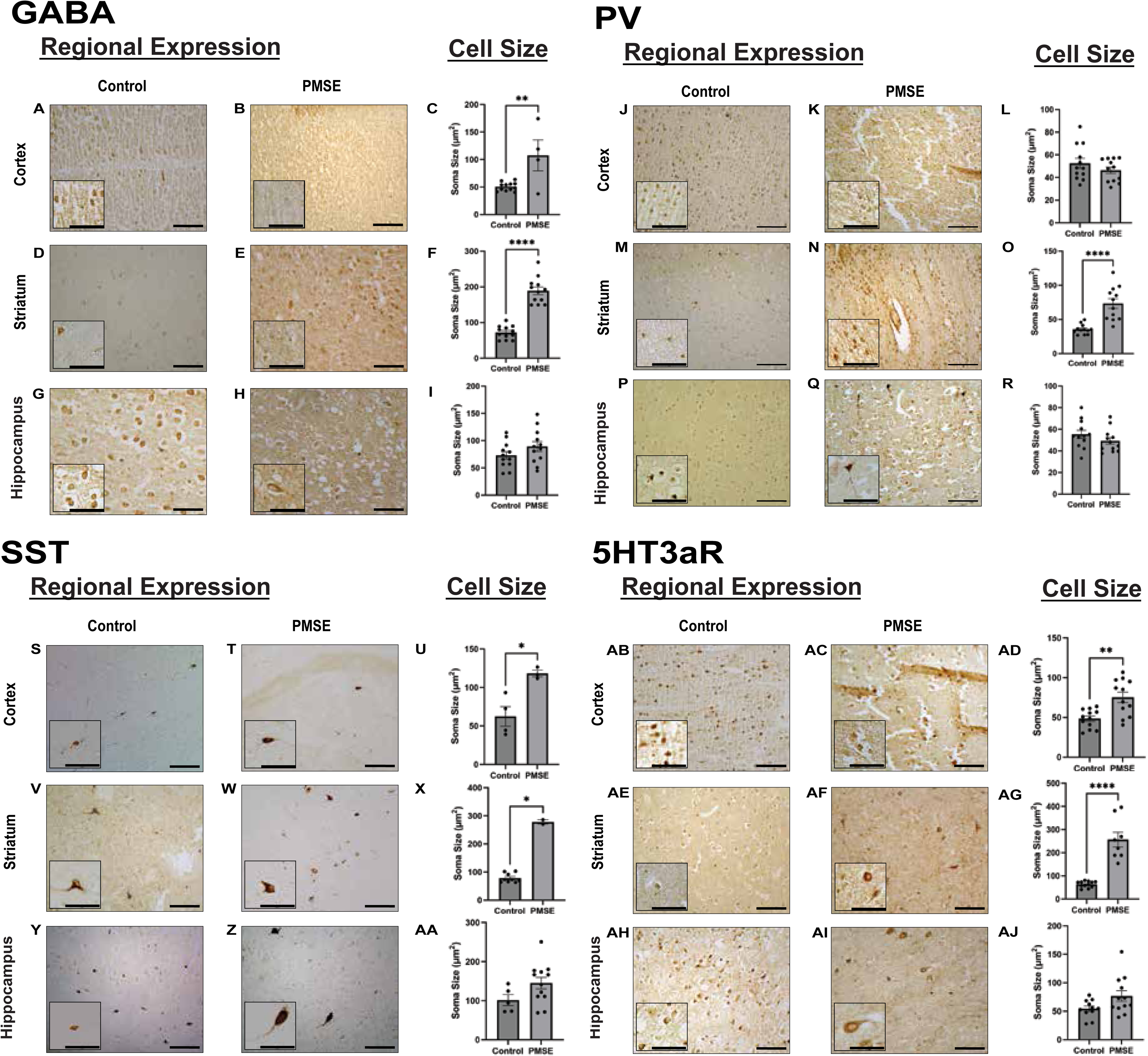
Immunohistochemical staining of human infant control and PMSE tissue. PMSE GABA+ interneurons (**A-I**) are decreased in number in the cortex and hippocampus and increased in the striatum, compared to control. This is replicated in the 5HT3aR+ interneurons (**AB-AJ**). PMSE PV+ interneurons (**J-R**) are decreased in number in the cortex and increased in the striatum. SST+ interneurons (**S-AA**) are decreased in number in the hippocampus and increased in the striatum. Cell size was increased in cortical GABA, 5HT3aR, and SST+ neurons, hippocampal GABA, PV, 5HT3aR, SST+ neurons but no striatal INs in PMSE tissue. Scale bars on larger images are 100µm, and 50µm on insets.

### PMSE INs exhibit region-specific cytomegaly

Cytomegaly is a known feature of mTOR hyperactivation in many cell types but has not been previously evaluated in PMSE INs.^1,3^ We quantified IN soma size across regions by tracing and measuring the soma of up to 10 representative cells in ImageJ. In cortex, soma size was increased in PMSE for GABA+, SST+, and 5HT3aR+ INs, and in hippocampus, GABA+, PV+, SST+, and 5HT3aR+ PMSE INs were enlarged. **(Figure 1, Supplemental Figure 1; Supplemental Table 3).** These data demonstrate IN cytomegaly in PMSE, suggesting that INs, like their excitatory counterparts, are enlarged likely due to STRADA loss and mTOR hyperactivation.^8^

### *Strada^-/-^* mice exhibit reduced cortical INs, increased striatal INs, and cytomegalic striatal INs at P21

Immunolabeling for major (PV, SST, and 5HT3aR) and minor (calbindin [CB], neuropeptide Y [NPY], and calretinin [CR]) IN subtypes was performed in cortex and striatum at P21. Cell counts were compared across WT, Het, and KO animals to assess genotype-dependent differences in regional expression consistent with altered migration.

In cortex, *Strada^-/-^* mice exhibited *reduced* numbers of GABA+ and PV+ cells and *increased* numbers of NPY+ cells compared with WT and Het animals. GABA+ cortical interneurons were enlarged in the *Strada^-/-^*mice, and their loss was biased towards cortical layers 2-3. **(Fig 2).** Laminar analysis was performed for all major subtypes but revealed no significant differences, suggesting that the changes in numbers of GABAergic subtypes was uniform across all cortical layers. In striatum, *Strada^-/-^* mice exhibited *increased* numbers of GABA+, PV+, SST+, and CB+ cells, with a concomitant decrease in 5HT3aR+ cells. No differences were observed in Lim homeobox 6 (Lhx6)+ progenitors across genotypes in either region, indicating that changes in IN numbers were likely not explained by a failure of IN progenitor cells to differentiate (see **Supplemental Table 3).** Cortical IN loss with striatal accumulation in the *Strada^-/-^* mouse provides solid statistical evidence to support similar findings in a single PMSE human tissue specimen and supports a Strada-associated IN migratory defect, with particular impact on PV+ INs. In sum, these observations suggest an enhanced effect of Strada loss on early MGE-derived IN migration. Additionally, GABA, PV, SST, and NPY labeled cells were enlarged in the striatum in *Strada^-/-^* mice with a trends towards CB+ IN enlargement (p=.0537), suggesting a phenotype of mTOR-hyperactivity in MGE-derived interneurons that have failed to migrate (**Figure 3**). This was further corroborated by increased P-S6 intensity in the striatum of P21 *Strada^-/-^*mice to determine if mTOR hyperactivity was playing a role in striatal retention (**Supplemental Figure 4**). Of note, we analyzed GABA and PV expression patterns across multiple hippocampal subregions and found no significant differences between WT, Het, and *Strada* KO (**Supplemental Figure 6**).

**Figure 2:**
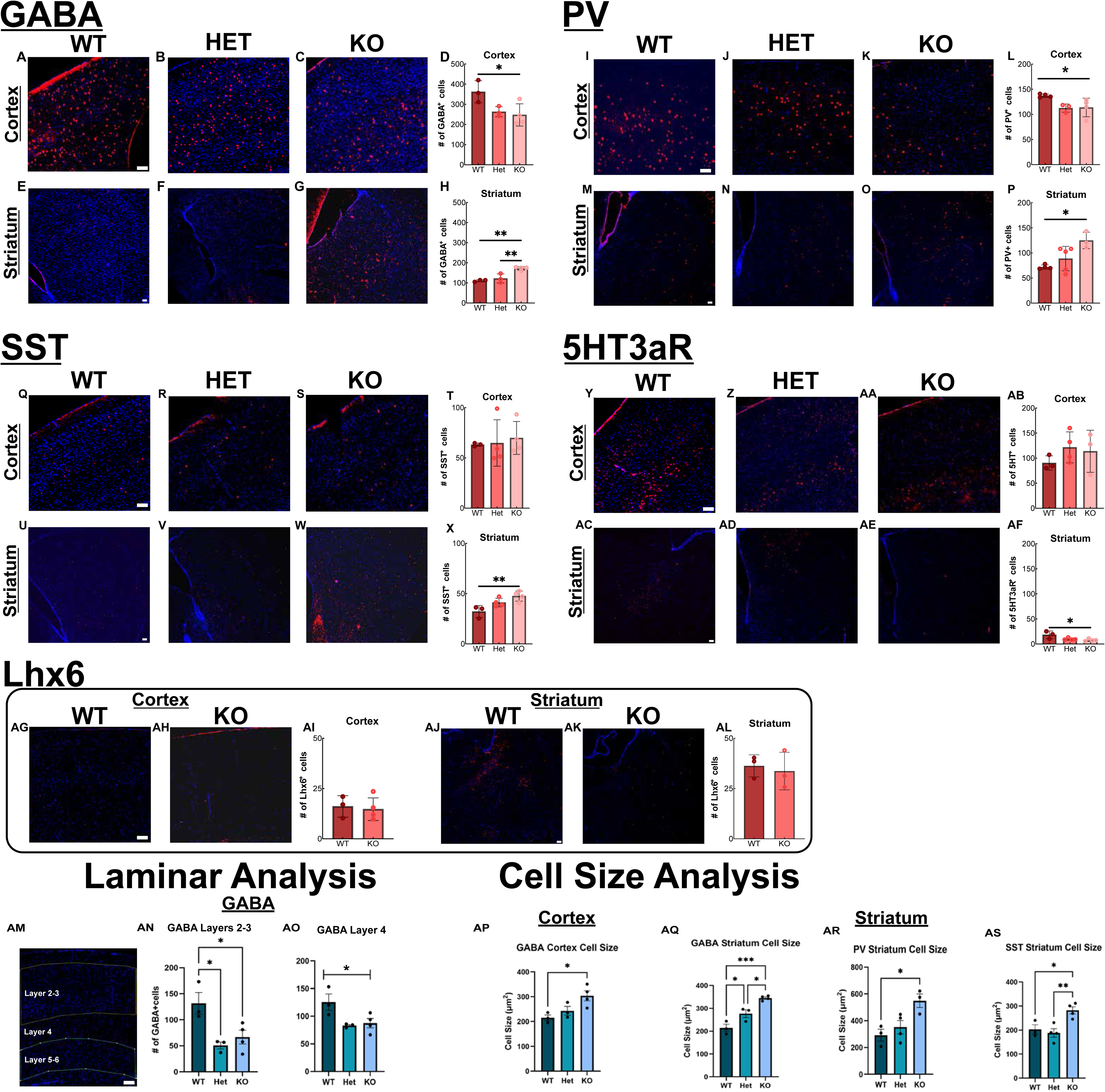
Immunofluorescent staining of major interneuron subtypes in P21 mouse WT, Het, and *Strada* KO cortical and striatal tissue. Strada KO cortex has decreased number of GABA+ (**A-D**) and PV+ (**I-L**) neurons with no change in SST+ (**Q-T**) or 5HT3aR+ (**Y-AB**), compared to control. KO striatum has increased numbers of GABA+ (**E-H**), PV+ (**M-P**), and SST+ (**U-X**) neurons and decreased numbers of 5HT3aR+ neurons (**AC-AF**), compared to control. There is no change in Lhx6 labeled cells (**AG-AL**) across genotype and brain region. Laminar analysis, based on DAPI density revealed a decrease in the number of GABAergic INs in layers 2-3 and layer 4 of the KO cortex compared to control (**AM-AO**). Soma size was increased in cortical GABAergic INs, and striatal GABAergic, PV, and SST+ INs in the KO mouse (**AP-AS**). Scale bars are 100µm. *p<.05, **p<.01, ***p<.001, ****p<.0001.

**Figure 3:**
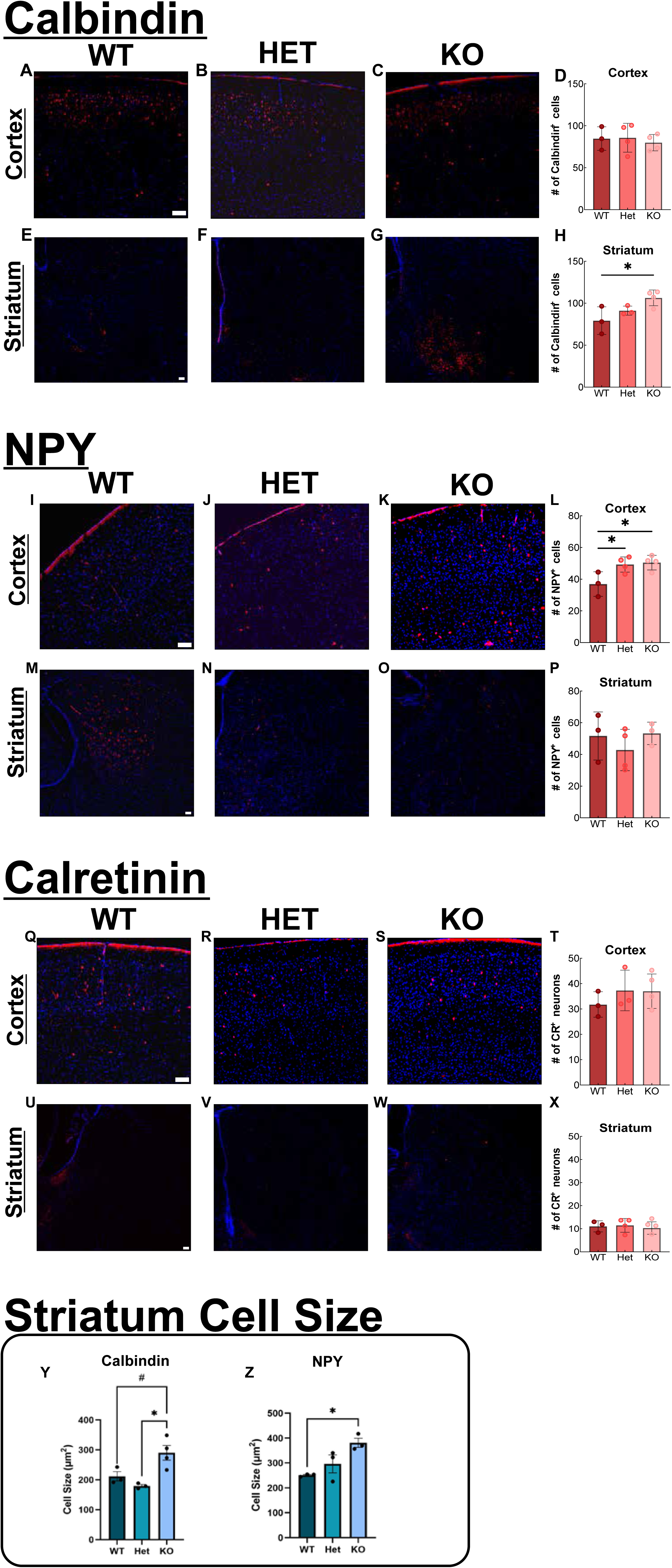
Immunofluorescent staining of minor interneuron subtypes in P21 mouse WT, Het, and *Strada* KO cortical and striatal tissue. KO cortex has increased numbers of NPY+ neurons (**I-L**) and no change in numbers of CB+ (**A-D**) and CR+ (**Q-T**) neurons, compared to control. KO striatum has increased numbers of CB+ neurons (**E-H**) and no change in numbers of NPY+ (**M-P**) or CR+ (**U-X**) neurons, compared to control. Soma size was increased in CB and NPY+ (**Y-Z**). Scale bars are 100µm. *p<.05, **p<.01, ***p<.001, ****p<.0001, #p=.0537.

### Strada loss alters gene expression related to IN development, migration, cytoskeletal organization, and mTOR signaling

To further investigate the developmental timing and regional effects of Strada loss on INs and the cytoskeleton, bulk RNAseq was performed on WT and *Strada^-/-^* P3 mouse cortex and P21 mouse cortex, hippocampus, and subcortical regions (basal ganglia and thalamus, representing remnant GE). Differentially expressed genes (DEGs) between WT and KO were identified at p<0.05. Prior to analysis, a prospective curated set of 193 functionally relevant genes, chosen based on functions related to IN development, cell migration, cytoskeletal organization, and mTOR signaling was created and used for regional analysis.

In P3 cortex, 74/193 genes of interest were altered in KO (35 upregulated, 39 downregulated). In P21 cortex, 31/193 genes were altered (20 upregulated, 11 downregulated). In P21 subcortex, 82/193 genes were altered, (62 upregulated, 20 downregulated), including multiple interneuron-associated genes transcripts, such as *Dlx2*, *Vip*, *Dlx5*, and *Sst*. In P21 hippocampus, 9/193 genes were altered (6 upregulated, 3 downregulated). *Strada* mRNA expression was significantly reduced in all KO regions. The full list of significant (p<0.05) DEGs per brain region is available in **Supplemental Table 5.**

Functional category analysis revealed *Fezf2* to be the most upregulated IN-associated transcript in P21 KO subcortex, and *Pvalb* to be the most down regulated in P3 cortex. *Satb2*, *Neurod2,* and *Sp8* were among the most upregulated migration-related genes in P21 subcortex. *Neurod6* was the most upregulated cytoskeletal gene in P21 subcortex, while *Tpp3* was the most downregulated in P3 cortex. Among mTOR-associated genes, *Creb5* was the most upregulated and *Lamp5* was the most downregulated in P3 cortex. Our transcriptomic findings demonstrate significant effects of Strada loss in the developing cortex and developed striatum. Of note, although all regions exhibited dysregulation across functional categories, DEGs were most abundant in the P3 cortex and P21 subcortex, suggesting an early developmental mechanism and ongoing dysregulation in remnant germinal regions **(Figure 4; Supplemental Table 4, Supplemental Figure 6)**.

**Figure 4:**
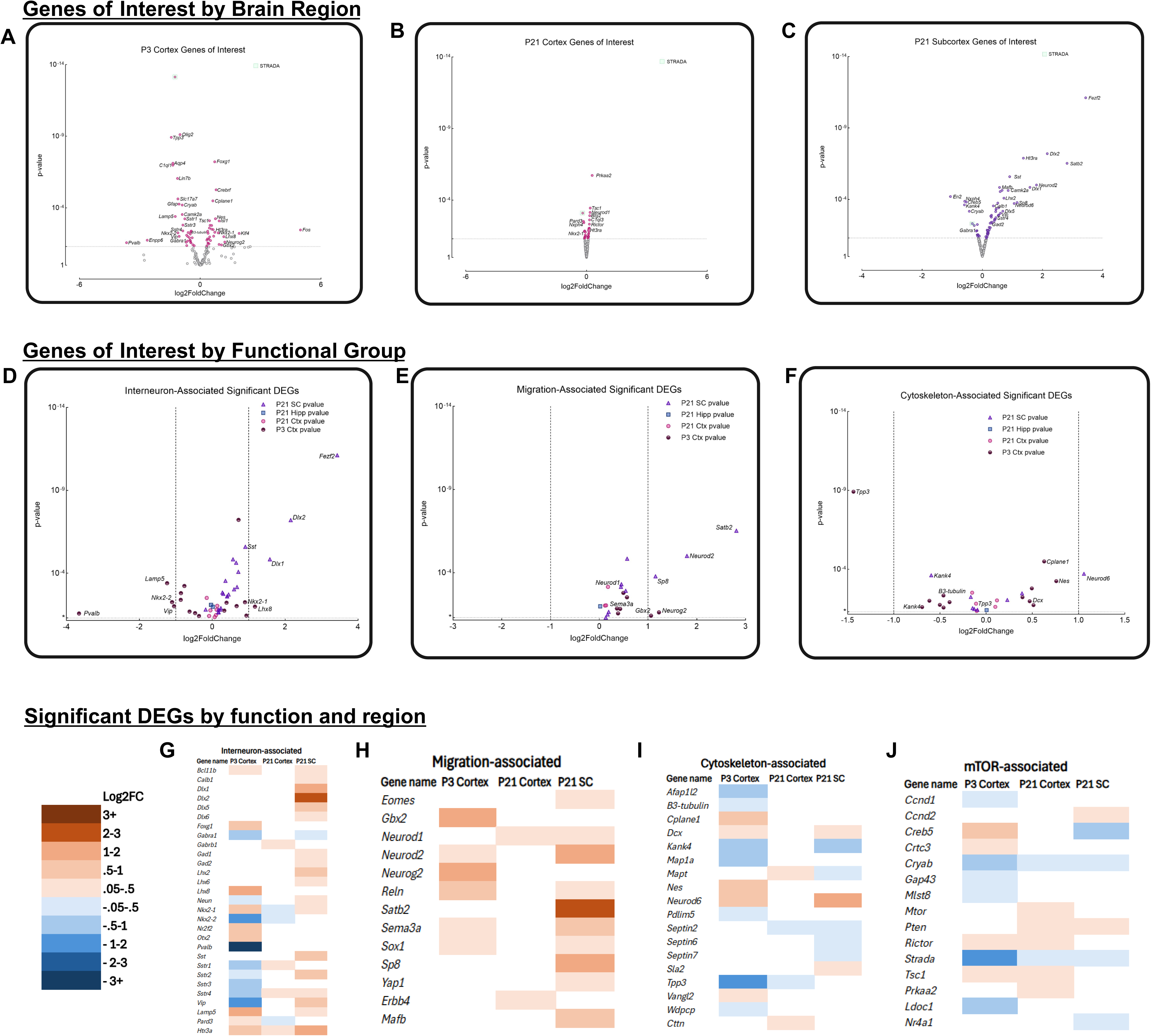
Bulk RNA-seq analysis of P21 and P3 WT and *Strada* KO mouse brain regions. Significant DEGs (p<0.05) from our genes of interest by brain region (**A-C**) and functional group (**D-F**) are represented, with regions and groups as labeled and discussed in text. **G-J**: Heat map depicting upregulated and downregulated DEGs (log2foldchange), classified by functional group and brain region.

## DISCUSSION

Multimodal analysis of human PMSE and age-matched control brain and *Strada* KO, heterozygous, and WT mouse brain demonstrates that STRADA/Strada loss impairs cortical GABAergic IN development and lamination. Histological and transcriptomic analyses in mice support a model in which Strada loss disrupts IN development by impairing tangential migration from the GE to the cortex and hippocampus, with pronounced effects on MGE-derived subpopulations. To our knowledge, this is the first study implicating IN migration impairment in the developmental pathogenesis of an mTOR-associated megalencephaly syndrome.

In mice, inhibitory NPC tangential migration begins at embryonic day (E) 11.5, peaks around E14, and continues through P10, corresponding to the second trimester through early postnatal development in humans.^12,16,21^ Our histological findings revealed that Strada loss was associated with reduced cortical INs and a corresponding accumulation in the striatum, a remnant of the developed GE.^22^ Interestingly we found a similar alteration in INs in a single PMSE brain specimen. Importantly, neural progenitor numbers were unchanged, indicating that cortical IN loss was unlikely to be due to impaired progenitor generation or differentiation.

### Strada Loss Causes Altered IN Migration

*Strada*^-/-^ mice exhibit histopathological IN changes that closely parallel those of human PMSE. Cortical loss of GABA+ and PV+ neurons with increased striatal GABA+, PV+, and SST+ populations supports a conserved defect in MGE-derived IN migration. The loss of GABA+ INs biased towards cortical layers 2-3 parallels our previous finding in excitatory cortical neurons that *Strada* loss causes an accumulation of upper-layer fated cells (Cux1+) in deeper cortical layers, perhaps in an mTOR dependent manner, suggested by the cytomegaly of cortical GABA+ INs in the P21 KO mouse.^8^ Although the number of 5HT3aR+ cells was unchanged in the KO cortex, a decrease in 5HT3aR+ cells in the striatum suggests differential effects on MGE- versus CGE-derived populations in mice. Species-specific differences in the proportion of 5HT3aR+ interneurons and general cortical inhibitory neuron composition potentially contribute to disparities between mouse and human phenotypes.^23^ Overall, the consistent reduction of cortical INs and accumulation of striatal INs across major subtypes in both *Strada* KO mice and PMSE tissue supports a Strada-associated migratory failure of inhibitory NPCs during neurodevelopment.

To distinguish migratory failure from impaired differentiation, we quantified Lhx6+ neurons, a marker of immature MGE-derived interneurons, directly downstream of Nkx2.1.^24^ If differentiation were disrupted, an accumulation of Lhx6+ cells would be expected in cortex or subcortex. However, unchanged Lhx6+ cell numbers in cortex and striatum between KO and WT mice indicate preserved differentiation capacity and support a primary migratory defect for INs.

Analysis of minor interneuron subtypes further suggests subtype-specific effects and compensatory mechanisms. For example, CB+ neurons, a minor SST+ IN subtype, are increased in striatum, while NPY+ cells are increased selectively in KO cortex, suggesting that the effects of Strada loss might differ between minor subtypes of interneurons.^25,26^ In contrast, CR+ interneurons, which are largely CGE-derived, are unchanged in both cortex and striatum.^27^ Together with preserved cortical 5HT3aR+ (CGE-derived) populations, these findings suggest that MGE-derived INs may be more vulnerable than CGE-derived populations to Strada loss. Whether this larger impact of Strada loss on MGE versus CGE progenitor populations reflects an anatomical regional signaling difference or an enhanced role of Strada at different neurodevelopmental timepoints, as MGE and CGE progenitor migration patterns vary temporally, remains to be determined.^9,28^

Disruption of PV+ cortical INs has been previously described in association with mTOR dysregulation in TSC, with evidence of a developmental phenotypic switch between PV-fated and SST-fated interneurons.^29^ PV+ INs have also been well-characterized in epilepsy, intellectual disability (ID), autism-spectrum disorder, Dravet Syndrome, and several other neurological disorders.^30,31^ These interneurons synapse with both pyramidal cells and other interneurons and tightly regulate neuronal oscillations and excitability^31^. Thus, a significant loss of PV+ INs in PMSE could directly contribute to the fully penetrant phenotypes of ID and intractable epilepsy.

Both cytomegaly of MGE-derived neurons and mTOR hyperactivation (elevated P-S6) in the striatum suggests that a migration impairment of inhibitory NPCs from the striatum to the cortex may be caused by mTOR hyperactivation and cytomegaly. Cytomegaly, as seen in other mTORopathies such as FCD Type IIB, is associated with aberrant actin and microtubule polymerization, which also govern the migration process.^32^ More work is needed *in vitro* to test the impact of Strada on the cytoskeletal dynamics of migrating NPCs.

### IN cortical and hippocampal deficits and striatal retention in human PMSE brain

While we acknowledge that a single PMSE brain specimen is not an adequate sample size to form solid conclusions about INs in PMSE, we submit that the similarities between findings in Strada-/- mice and human PMSE are compelling. Reductions of GABA+, GAD65+, PV+, and 5HT3aR+ INs in PMSE cortex and reductions of GABA+, SST+, and 5HT3aR+ INs in PMSE hippocampus compared with controls as well as *increased* numbers of INs across all major subtypes in striatum suggests that the developmental effects of STRADA loss in humans are not restricted to a single IN lineage. Because the GE gives rise to subcortical structures such as the basal ganglia, including striatum, and thalamus, retention of INs in this region is consistent with migratory failure.^22^ Notably, there is no difference in the number of NKX2.1+ MGE-derived progenitor cells across cortex, hippocampus, or striatum in PMSE versus control tissue. Together, the accumulation of interneurons in the striatum at the expense of cortical and hippocampal populations, in the absence of progenitor deficits, supports impaired migration of inhibitory NPCs from the GE to their laminar destinations. Impaired cytoskeletal regulation, previously demonstrated in Strada-deficient excitatory NPCs, may underlie this migratory failure, though direct testing in inhibitory NPCs is needed.^7^

As in the Strada-/- mouse we further identified IN cytomegaly in PMSE tissue, extending prior observations of cytomegaly in other neuronal populations in PMSE. Cytomegaly serves as a key feature of mTOR dysregulation in neurons across multiple mTORopathies, including TSC and Focal Cortical Dysplasia (FCD) type IIB.^4,6,33^ Our work represents the first report of IN cytomegaly in an mTOR-associated megalencephaly and suggests that mTOR hyperactivity resulting from STRADA loss substantially affects GABAergic IN populations

### Developmentally timed alterations in PMSE gene expression

RNAseq analysis identified numerous genotype-, region-, and age-dependent transcriptional differences. Our genes of interest had the most robust expression differences in *Strada* KO in the P3 cortex and P21 subcortex (basal ganglia and thalamus) across all 4 functional categories. In the P3 KO cortex, interneuron-, cytoskeleton-, and mTOR-related genes were predominantly downregulated, including cytoskeletal regulators (*Tpp3*, *Map1a*, *Tubb3*, *Mbp*), IN markers and receptors of each major IN subtype (*Pvalb*, *Vip*, *Sstr1-4*, *Gabra1*), and the mTOR pathway gene *Mlst8*. In contrast, migration-associated genes (*Sema3a*, *Reln*), regional markers (*Nkx2-1*, *Nkx2-*2, *Lhx8*, *Foxg1*, *Gbx2*), mTOR-related gene *Rictor*, and immediate early genes (*Fos*, *Fosb*) were upregulated. Functional validation will be needed to determine whether these transcriptional changes are consistent with disrupted IN migration and lamination in early brain development.

At P21, fewer transcriptional differences were observed in cortex and hippocampus, consistent with the lack of observed differences in the number of inhibitory neurons in the P21 hippocampus of *Strada*^-/-^ mice **(Supplemental Figure 6)**. Some cytoskeletal genes (*Tpp3*, *Mbp*) remained downregulated in the P21 KO cortex, but unlike the P3 cortex, regional markers *Nkx2-1* and *Nkx2-2* are downregulated in the P21 KO cortex. *Htr3a* and *Fos* continue to be upregulated in the P21 cortex, along with interneuron receptor genes (*Sstr1*, *Sstr4*, *Gabrb1*), several of which were downregulated in the P3 cortex. This may be due to compensatory mechanisms to counteract the loss of interneurons.^34^ Several mTOR pathway genes were also upregulated (*Mtor*, *Rictor*, *Ulk1*, *Tsc1*, *Pten*), which may contribute to cortical hyperexcitability, consistent with prior observations in Strada-deficient excitatory neurons.^8,35^ In the P3 KO cortex, there is an alteration of the expression of several migration and regional markers of interneurons, many of which are unchanged in the P21 KO cortex, suggesting that the processes regulated by these genes are most strongly affected in the early developing brain. This parallels the timeline of neural progenitor migration, which is tightly regulated by brain region and migratory cues, many of which have muted effects after completion of corticogenesis. In the developed brain, dysregulation of mTOR- and cytoskeleton-associated genes in the cortex can likely directly impact both excitatory and inhibitory neuronal activity, contributing to the epilepsy phenotype that persists in PMSE.^36^

The P21 subcortex showed extensive upregulation of IN and progenitor markers including *Nkx2-1, Lhx6, Gad1/2, Dlx* family members, *Sst, Vip*, and *Npy*, consistent with an IN accumulation in this region. Immediate early genes (*Fos*, *Fosb*, *Npas4*) were also upregulated, suggesting increased neuronal activity associated with excess subcortical neurons.^37–39^ Additionally, several cytoskeleton-associated transcripts are downregulated, including *Sep2*, *Sep6*, *Sep7*, *Kank4*, and *Mbp.* These findings, paired with a downregulation of regional and cytoskeletal-related transcripts in the P3 cortex, further support impaired migration driven by disrupted cytoskeletal dynamics.

In the cell identity-associated transcripts across all brain regions, the most upregulated genes were in the P21 *Strada* KO mouse cortex, with the transcripts *Fezf2*, *Dlx1*, and *Dlx2* having the highest fold changes. *Dlx1* and *Dlx2* are present in multiple classes of developed INs with a preference for SST+ cells, and *Fezf2* is a marker of layer 5 cortical projection neurons^40–42^. The highest number of cytoskeleton-related genes were dysregulated in the P3 cortex, with *Tpp3* being the most downregulated. Other dysregulated genes include *B3-tubulin*, *Map1a*, and *Dcx*. Widespread dysregulation of cytoskeletal transcripts, particularly in the P3 cortex, supports impaired migration potentially driven by disrupted cytoskeletal dynamics.^21^

### Limitations

Low neonatal viability in *Strada^-/-^* mice limited histological analysis to a single postnatal time point and precluded the ability to determine the mTOR-dependence of our findings with a rapamycin treatment trial. Although P21 precedes full synaptic maturation, prior work and our transcriptomic data indicate that Strada-associated pathology is primarily developmental, and all PMSE patients start experiencing symptoms of the disease, including epilepsy, at or within a few months of birth.^1,43^ Of note, ours is the first study of the adult *Strada^-/-^* mouse brain.

Survivorship bias may influence *Strada* KO mouse analyses due to high observed rate of pre- and perinatal lethality, which is consistent with observations in *Lkb1* mutated mouse lines and with a high rate of miscarriage in mothers carrying fetuses with STRADA deficiency.^8,17,44^ Future studies investigating Strada loss in a focal KO model may be needed to mitigate this effect.

Additionally, human tissue availability is limited to a single PMSE brain, represented in our human histological analysis **(Figure 1)**, which is also the same brain tissue used in previous histological studies of PMSE.^1,3^ Introduction of error from extrapolation of results based on this n of 1 is mitigated by the fact that the represented PMSE *STRADA* mutation is consistent with the most commonly reported pathogenic disease-associated mutation (deletion of *STRADA/LYK5* exons 9-13), and PMSE is characterized by homogenous disease characteristics across the affected patient population^1^. Furthermore, we note remarkable consistency with IN and IN subtype regional expression patterns in the *Strada* KO mouse brains, suggesting that our human tissue results are less likely to be an outlier. Future studies will include any other tissue obtained in donation from PMSE patients.

Finally, this is a descriptive study based on histopathology and transcriptomics. Nevertheless, by leveraging complementary human and mouse models, it establishes a solid foundation for future mechanistic and interventional IN studies, including developmentally timed interventions such as mTOR inhibitors shown in previous work to mitigate the effects of Strada loss in mouse excitatory neurons.^7^

## CONCLUSION

In summary, this study provides the first evidence that STRADA/Strada deficiency disrupts inhibitory neuron lamination and induces IN cytomegaly in an mTOR-associated megalencephaly. Cortical IN loss with subcortical accumulation, in the absence of differentiation deficits, supports a primary migratory failure. Transcriptomic data further implicate early disruption of migration- and cytoskeleton-associated pathways. Ultimately, our discovery of developmentally disrupted IN migration and lamination with STRADA/Strada deficiency emphasizes the need for earlier therapeutic intervention in PMSE and perhaps other mTOR-associated megalencephalies, in order to modify aberrant IN cortical lamination and the loss of excitatory:inhibitory balance that likely contributes to key disease features such as epilepsy and neurocognitive impairment.

## ACKNOWLEDGEMENTS

Grant Support:

NINDS K08NS140393, TEDCO MSCRF grant to W.E.P.

We acknowledge Marianna Baybis, Janice Babus, Irina Sbornova, Aria Moss, David Kolb, and John Page for their thoughtful insights in discussion and interpretation of results. We acknowledge the Institute for Genome Sciences Core at the University of Maryland, Baltimore, specifically Carrie McCracken and Luke Tallon, for tissue processing and analysis for the RNAseq experiments and for assistance with data interpretation.

## AUTHOR CONTRIBUTIONS

R.K.P. contributed to conceptual design, experimental planning and implementation, data analysis and interpretation, and manuscript writing.

A.H. contributed to experimental planning and implementation and data analysis.

T.H.N. contributed to conceptual design and experimental planning and implementation.

M.P. contributed to experimental planning and implementation and manuscript construction.

R.Y-M. contributed to experimental implementation, data analysis, and manuscript construction.

D.A.A. contributed to experimental implementation and manuscript construction.

S.K. contributed to experimental planning and implementation.

S.T. contributed to data analysis

V.J.C. contributed to clinical interpretation, manuscript writing, and provision of PMSE tissue.

P.H.I. contributed to data interpretation and manuscript writing.

L.T.D. contributed to experimental planning, data interpretation, and manuscript writing.

P.B.C. provided the Strada-/- model, contributed to experimental planning and data interpretation and manuscript writing.

W.E.P. contributed to scientific concept, experimental planning, data analysis and interpretation, and manuscript writing.

