## Supplemental Materials for "STRADA deficiency impairs cortical interneuron development in humans and mice"

**SUPPLEMENTAL METHODS**

**Animals**

Between 3-5 WT (*Strada^+/+^*), heterozygous (Het, *Strada^+/-^*), and KO (*Strada^-/-^*) P21 male and female mice were used per group for histopathology. Three P3 WT and KO mice were used per group for bulk RNA-sequencing (RNAseq). Procedures were performed as described and were approved by the University of Maryland, Baltimore Institutional Animal Care and Use Committee under protocol AUP-00000032 to W.E.P.

The germline *Strada*^-/-^ mouse line was generated in the Crino lab^8^ using a neomycin cassette to model the murine equivalent most abundant Strada founder variant which was found the Old Order Mennonite community.^1,8^ This variant is a homozygous deletion of exons 9-13 of the *STRADA/LYK5* gene (17q23.3) in humans. Our mouse model replicates this mutation in the murine chromosome 11 *Strada* ortholog, *2610019A05Rik*. As previously reported, the *Strada*^-/-^ mice are megalencephalic, and their body-lengths are shorter than those of their WT and Het counterparts^2^. The *Strada* mutation in mice is associated with high levels of pre- and perinatal lethality^2,3^. PCR of sectioned, adjacent slides to those chosen for histology was performed to verify genotype, originally obtained by PCR of tail snips. (**Supplemental Figure 3**). To confirm well-characterized findings of mTOR hyperactivity in Strada loss, we immunolabeled its downstream target phospho(p)-S6^Ser240/244^ in the cortex and of WT and KO mice, and found increased expression in the KO brain (**Supplementary Figure 4**).^1,3,7,8^

**Histopathology**

*Tissue processing and preparation*

At P21, animals were euthanized by CO_2_ exposure and perfusion with ice-cold 1x phosphate buffered saline (PBS) (Quality Biological, #119-069-131) and 4% paraformaldehyde (PFA) (Electron Microscopy Sciences, #15710). Brains were extracted and immersion-fixed at 4°C in 4% PFA for 24 hours prior to processing. To begin processing, brains were incubated at 4°C in PBS overnight, processed in a series of ethanol dilutions increasing in concentration (70%, 80%, 90%, 95%, 100%) and xylene, before being embedded in paraffin. Brains were cut at room temperature into 10µm coronal sections on a sliding microtome and rehydrated in warm deionized water before being mounted onto charged slides. Slides were dried overnight in an oven set to 53°C and cooled at room temperature prior to immunohistochemistry.

*Immunohistochemistry: Mouse tissue*

PV and GABA: Slides were deparaffinized in xylene, followed by a series of ethanol decreasing in concentration from 100% to 70%, and rinsed in water. Slides were placed in a humidified chamber, and sections were blocked for 1hr at room temperature with a solution containing 1x PBS, .3% Triton-X (Sigma-Aldrich, X100-100ML), and 5% normal goat serum (Cell Signaling Technology, #5425). Following blocking, sections were incubated with primary antibody at a concentration of 1:500 diluted in blocking solution overnight at 4°C. Following primary antibody, slides were washed with PBS and incubated in fluorophore-conjugated secondary antibody for 2 hours at room temperature. Sections were washed with PBS and coverslips were applied with Fluoroshield with DAPI (Sigma-Aldrich, F6057-20ML).

5HT3aR, SST, CB, NPY, CR, LHX6: Slides were deparaffinized in xylene followed by a series of ethanol decreasing in concentration from 100% to 70% and rinsed in water. Following deparaffinization, sections underwent heat-mediated antigen retrieval using a sodium citrate unmasking solution (Vector, H-3300-250) and incubated in the sodium citrate until the temperature was cool to the touch, around 30 minutes. Slides were placed in a humidified chamber, and sections were blocked for 1 hour at room temperature with a solution containing 1x PBS, .2% Triton-X, and 5% normal goat serum. Following blocking, sections were incubated with primary antibodies at specified concentrations, diluted in blocking solution overnight at 4°C (see **Supplementary Table 1** for antibody and dilution details). Following primary antibody, slides were washed with PBS and incubated in fluorophore-conjugated secondary antibody (**Supplementary Table 1**) for 2 hours at room temperature. Sections were washed with PBS and coverslips were applied with Fluoroshield with DAPI.

Additionally, fluorescent co-labeling of GAD65 and GABA was performed to confirm antibody accuracy of interneuron labeling (**Supplemental Figure 2**).

*Immunohistochemistry: Human tissue*

Pediatric control cortex, hippocampus, and subcortical tissue from 2 different subjects (ages 40 days and 84 days; cause of death: possible SIDS and asphyxia by food; PMI: 30hrs and 20hrs) was obtained from the National Institutes of Health (NIH) tissue repository at the University of Maryland, preserved in 10% formalin. Tissue was rinsed in 1x PBS, processed in an ethanol series increasing in concentration from 70% to 100% followed by xylene, and embedded in paraffin. Sections were cut on a sliding microtome at room temperature at a thickness of 7mm to replicate the sectioning of the PMSE tissue, cut previously.^1,4^ Methods for PMSE patient tissue sectioning are described in Orlova et al. 2010. Sections were rehydrated in a water bath, directly mounted onto slides, and dried overnight at 53°C.

Prior to deparaffinization, slides were warmed at 53°C for 1 hour. Cooled slides were deparaffinized in xylene, followed by a series of ethanol decreasing in concentration from 100% to 70%, and rinsed in water. Heat-mediated antigen retrieval using sodium citrate was performed as previously described. Sections were blocked in 30% hydrogen peroxide (H_2_O_2_) (Sigma Aldrich, H1009-500ML) diluted in methanol (Sigma Aldrich, M1775-1GA) for 20 minutes. Following blocking, sections were rinsed with DI water, washed in 1x tris-buffered saline (TBS) (Quality Biological, #351-086-131) for 5 minutes and placed in 2% fetal bovine serum (FBS) (VWR, #97068-075) diluted in TBS for 5 minutes prior to overnight incubation in primary antibody. For SST, 5HT3aR, GABA and GAD65 staining but not PV, the 2% FBS solution also contained .2% Tween 20 (Bio Rad, #1706531). The antibody concentrations are listed in **Supplementary Table 1**. After incubation in primary antibody, sections underwent TBS and 2% FBS +/- Tween washes, followed by a 1-hour incubation in biotinylated secondary antibody (specified in **Supplementary Table 1**). Both the primary and secondary antibodies were diluted in the 2% FBS solution. Washes also followed the secondary antibody incubation, and this was followed by a 1-hour incubation in a solution containing avidin, biotin, and horse-radish peroxidase (HRP) (Vector, PK-6100) diluted in the 2% FBS solution. Tissue then underwent the final set of washes before incubation in 3,3’-Diaminobenzidine (DAB) (Sigma Aldrich, D4293-50SET) solution for 15-20 minutes or until optimal precipitation was achieved. Slides were then rinsed in TBS before sections were dehydrated in an ethanol series increasing in concentration (70%-100%), followed by xylene, and coverslipped with Permount (Fisher, SP15-100). Slides were then dried overnight in the fume hood prior to imaging.

Cytomegaly is a known feature of mTOR hyperactivation in many cell types but has not been previously evaluated in PMSE INs.^1,3^ We quantified IN soma size across regions by tracing and measuring the soma of up to 10 representative cells in ImageJ. In cortex, soma size was increased in PMSE for GABA+, SST+, and 5HT3aR+ INs, and in hippocampus, GABA+, PV+, SST+, and 5HT3aR+ PMSE INs were enlarged. **(Figure 1, Supplemental Figure 1; Supplemental Table 3).** These data demonstrate IN cytomegaly in PMSE, suggesting that INs, like their excitatory counterparts, are directly affected by STRADA loss and mTOR hyperactivation.

**Fluorescence imaging and quantification**

*Microscopy*

For each marker analyzed, 2 sections per animal were imaged (10x for cortex, 4x striatum) and fluorescent-bright cells were hand-counted using ImageJ software.^18^

The regions of interest (ROIs) in mouse brain were defined as follows, using the Allen Brain Atlas of Mouse (Allen Institute, Seattle, WA): The cortex is selected to encompass the somatosensory cortical region dorsal to the hippocampus and lateral to the anterior cingulate cortex and spanning the pial surface to the corpus callosum. The striatum is selected as the area between the lateral ventricle and the ventral corpus callosum, representative of remnant GE.^21^ Of note, a comparative more rostral cortical region at the level of the striatum in the anterior-posterior axis was selected for comparison to more terminal regions of later IN migration in the rostral migratory stream (**Supplementary Figure #**). Images were taken on a Keyence BZ-X800 fluorescence microscope (Keyence Corp, Itasca, IL) and equally adjusted for black balance.

*Quantification*

For each animal, 3 images were taken at each ROI, bilaterally when available, with consistent microscope settings. Cells were hand-counted using ImageJ, and a positive cell was defined both by morphology and nuclear colocalization with DAPI. For each animal, hand counts were averaged per ROI, and each average was plotted as a single data point per animal. This was completed for each antibody stain, as shown in **Figures 2 and 3.**

For the laminar analysis, the cortex was divided into upper (layer 2-3), middle (layer 4), and lower (layer 5-6) thirds based on DAPI density and each section was counted and compared separately across genotypes using a one-way ANOVA with Tukey’s multiple comparisons. Cell size in the striatum was measured by tracing the soma of 10 representative cells per section and comparing the average soma size per animal across genotypes using a one-way ANOVA with Tukey’s multiple comparisons.

*Statistics*

All statistical significance was determined using a one-way ANOVA per interneuron marker per brain region, with Tukey’s multiple comparisons post-hoc when appropriate, using Prism (GraphPad Software, Boston, MA). Significance was determined by a p-value < 0.05.

**Brightfield imaging and quantification**

*Microscopy and Quantification*

Human tissue was imaged on a Keyence microscope at 20x and 40x, with consistent microscope settings and white balance. Differences between the PMSE brain and control brain tissue were determined by semi quantification in ImageJ. 20x images were converted to 16-bit images and set to an auto-threshold of 1%, mitigating background staining while preserving all antibody-positive cells. Particles were then automatically counted using a set radius of 75 pixels.

To measure cell size, images were thresholded in the same manner, and the area of each cell was measured using ImageJ. Control and PMSE cells were compared using a Student’s T-test with significance determined by a p-value less than 0.05.

**Bulk RNA sequencing**

*Tissue preparation*

At P21, 3 WT and 3 KO mice were euthanized by CO_2_ exposure, followed by confirmatory cervical dislocation. Brain tissue was immediately extracted, rinsed in ice-cold PBS, and the forebrain was dissected into cortical, hippocampal, and subcortical (inclusive of thalamus and basal ganglia, to include remnant GE) areas under microscopic visualization. A separate set of 3 WT and 3 KO mice were euthanized and brains extracted in a similar manner at P3, from which only cortex was dissected. Tissue was immediately frozen on dry ice and stored at –80°C prior to bulk RNA-sequencing at the Institute for Genome Sciences Core at the University of Maryland, Baltimore.

*Quality Control*

FastQC software was used to assess sequence quality of all RNA samples submitted from all animals, measuring total mapped reads, percent mapped for each sample, and base call rate, represented by a BoxWhisker plot. Higher percentage mapped and base call rates indicate higher sample quality with less chance of contamination. All samples used were of high quality, greater than the 90^th^ percentile across all measures (**Supplemental Figure 7**).

*Analysis*

Data was processed using nf-core/rnaseq v3.18.0 (doi: [10.5281/zenodo.1400710](https://doi.org/10.5281/zenodo.1400710)) of the nf-core collection of workflows ([Ewels](https://doi.org/10.1038/s41587-020-0439-x) *[et al.](https://doi.org/10.1038/s41587-020-0439-x)*[, 2020](https://doi.org/10.1038/s41587-020-0439-x)), utilizing reproducible software environments from the Bioconda ([Grüning](https://doi.org/10.1038/s41592-018-0046-7) *[et al.](https://doi.org/10.1038/s41592-018-0046-7)*[, 2018](https://doi.org/10.1038/s41592-018-0046-7)) and Biocontainers ([da Veiga Leprevost *et al.*, 2017](https://doi.org/10.1093/bioinformatics/btx192)) projects.

The pipeline was executed with Nextflow v24.10.4 ([Di Tommaso *et al.*, 2017](https://doi.org/10.1038/nbt.3820)) with the following command: *nextflow run /local/projects/grc/devel/packages/nf-core-rnaseq_ 3.18.0/3_18_0/ --input /local/projects-t3/XPARK/process_log/GRC-6931/sample_sheet.csv --outdir /local/projects-t3/XPARK/process_log/GRC-6931 /ANALYSIS/RNASEQ_rsem_ 20250813_101842 --aligner star_rsem –gtf /local/db/transcriptomics_references/ m_musculus/embl/Mus_musculus.GRCm39.113.gtf--- fasta /local/db/ transcriptomics_references/m_musculus/embl /Mus_musculus. GRCm39.dna.toplevel.fa -c custom_slurm.config -profile singularity --multiqc_config /local/projects-t3/XPARK/ process_log/GRC-6931/multiqc_config_NEW.yaml --star_index /local/projects/grc/devel/ kvaviko/STAR_indexes/mouse.GRCm39.113 --salmon_index /local/projects/grc/devel/ kvaviko/SALMON_indexes/mouse.GRCm39.113 -resume*

Target genes of interest were isolated and categorized based on biological significance. Logarithmic changes and p-values were plotted by region of interest and functionality. The normalized gene count was used to compare expression of genes of interest in the cortex and subcortex (cortex/cortex+subcortex). Ratios of cortical to subcortical expression for KO mice were compared to WT mice via a two-tailed t-test. Calculations and statistics were done in Excel (Microsoft Office, 2025). For all statistics, significance was set at a p-value of 0.05.

Following bioinformatics analysis, differentially expressed genes (DEGs) were filtered to only represent transcripts with significant p-values for each brain region. Additionally, a list of 193 genes of interest was prospectively generated, to represent transcriptomic changes in particularly relevant functional pathway categories suspected to play a role in IN development in PMSE. These categories include: IN maturation, cell migration, cytoskeletal organization, and mTOR signaling. Analysis of these 193 genes was sub-divided by functional category and by brain region.

**SUPPLEMENTAL FIGURE LEGENDS**

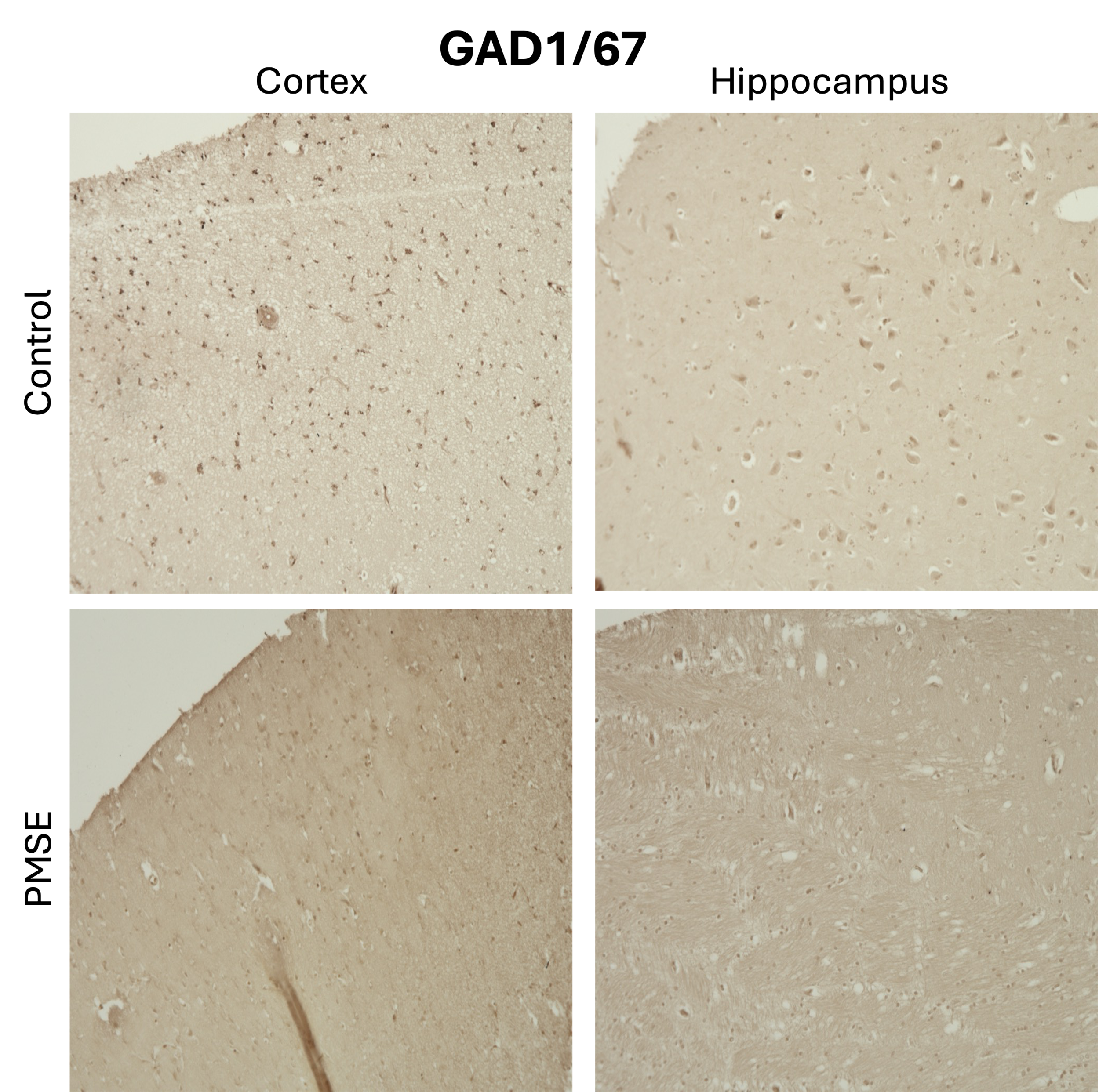

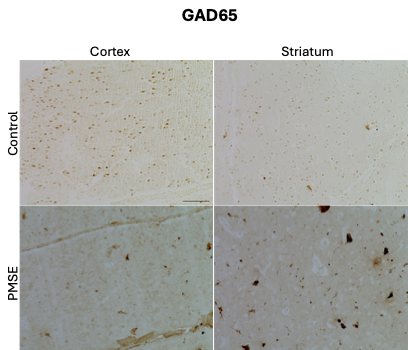

**Supplemental Figure 1**: GAD1/67 and GAD65 immunohistochemical labeling of control and PMSE tissue reveals a decrease in GAD65 and GAD67+ neurons in the PMSE cortex and an increase in GAD65+ neurons in the hippocampus. Images are taken at 20x.

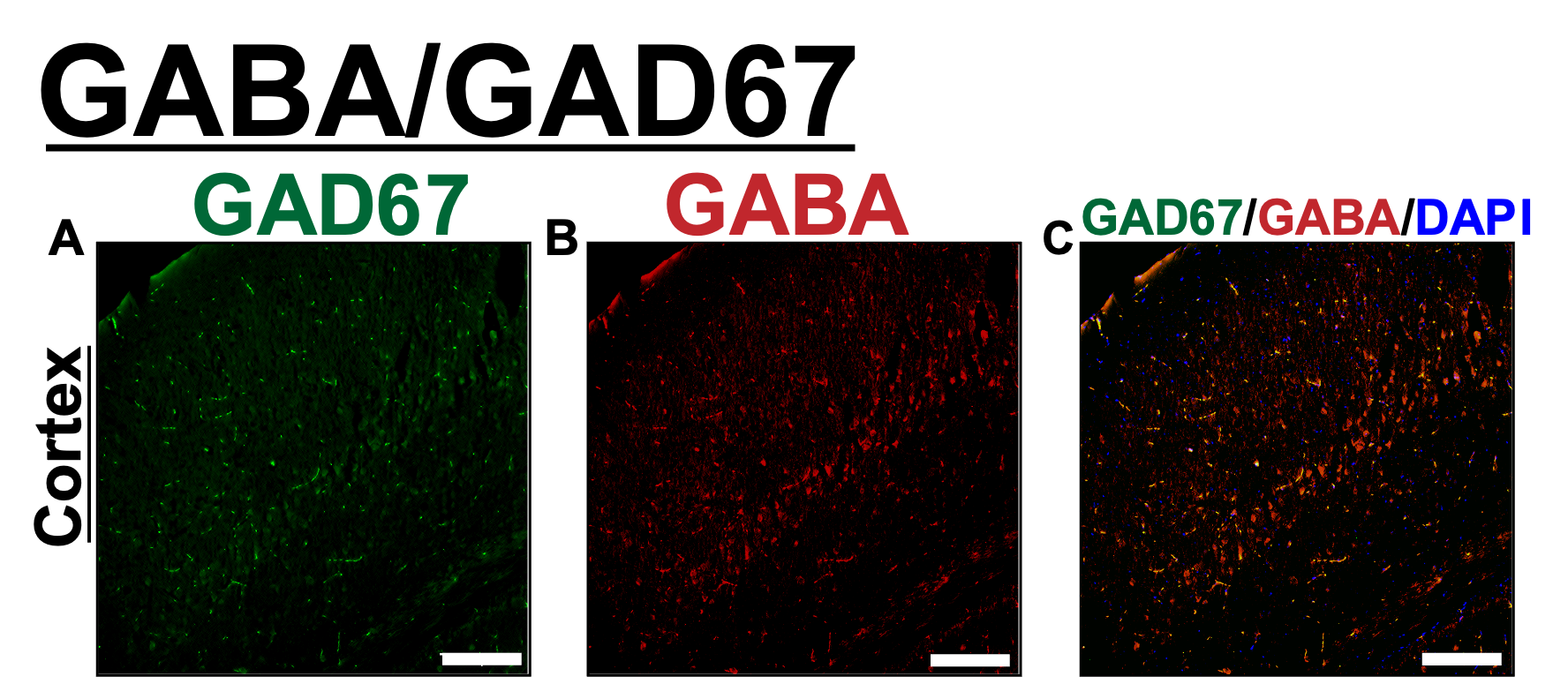

**Supplemental Figure 2**: Fluorescent co-labeling of GAD65 (**A**) and GABA (**B**) to ensure accurate quantification of interneurons. Images are taken at 10x. Scale bar is 100µm.

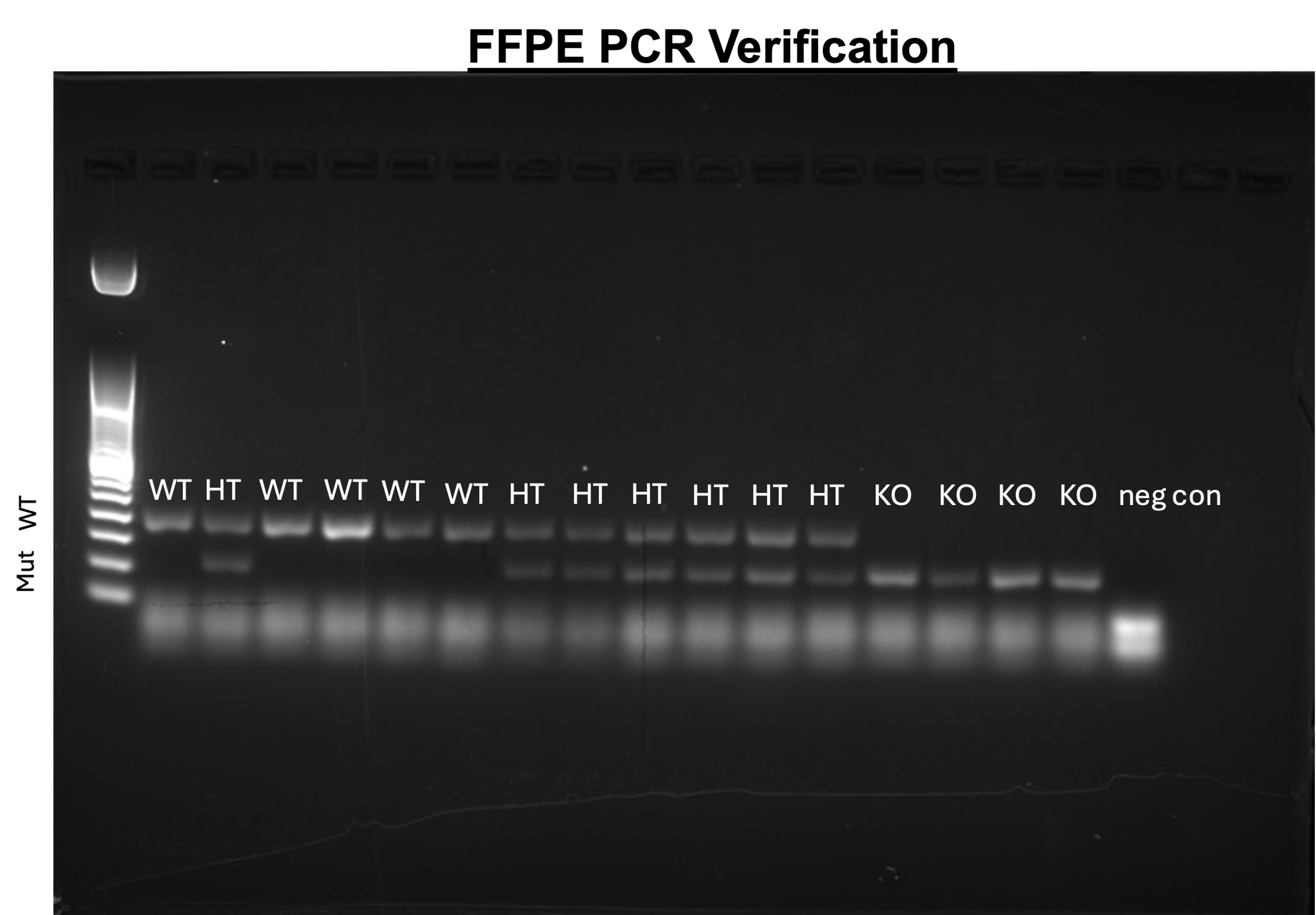

**Supplemental Figure 3:** Verification of *Strada* mouse genotypes used for histological analysis by polymerase chain reaction (PCR) of fixed and processed FFPE tissue on adjacent slides to those used for immunolabeling.

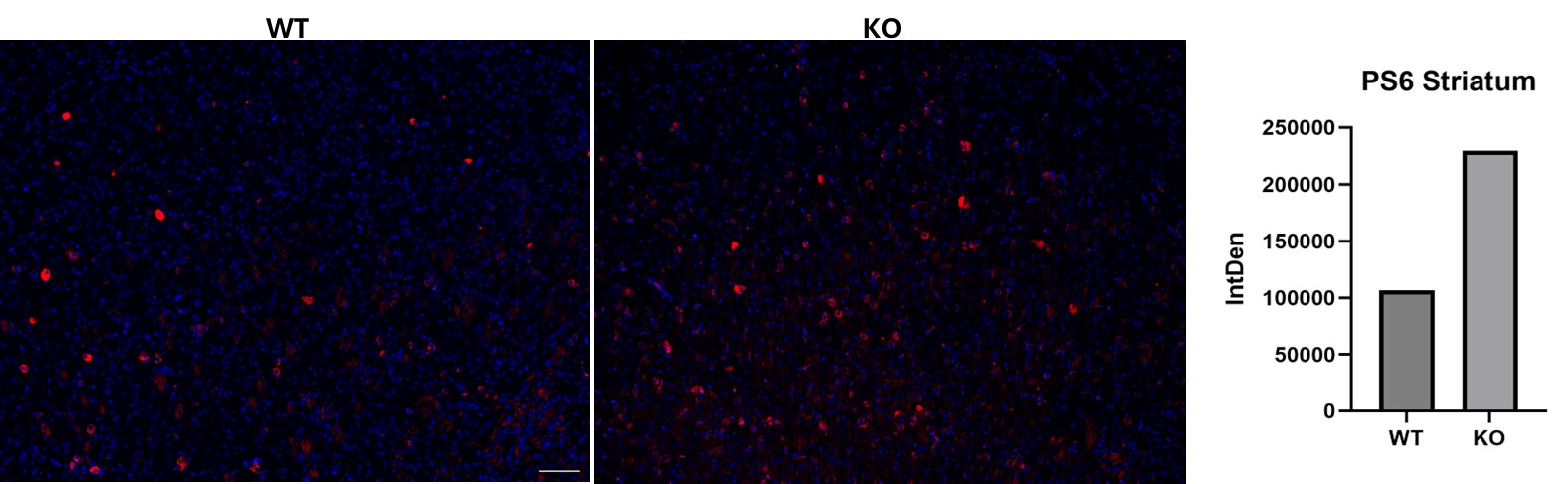
**Supplemental Figure 4:** Immunolabeling of elevated phospho-S6^240/244^ (PS6) in P21 KO striatum compared to WT to functionally validate mTOR hyperactivity previously observed with *Strada* loss, specifically in striatal remnant cells as a potential mechanism of failed migration. Scale bars are 100µm.

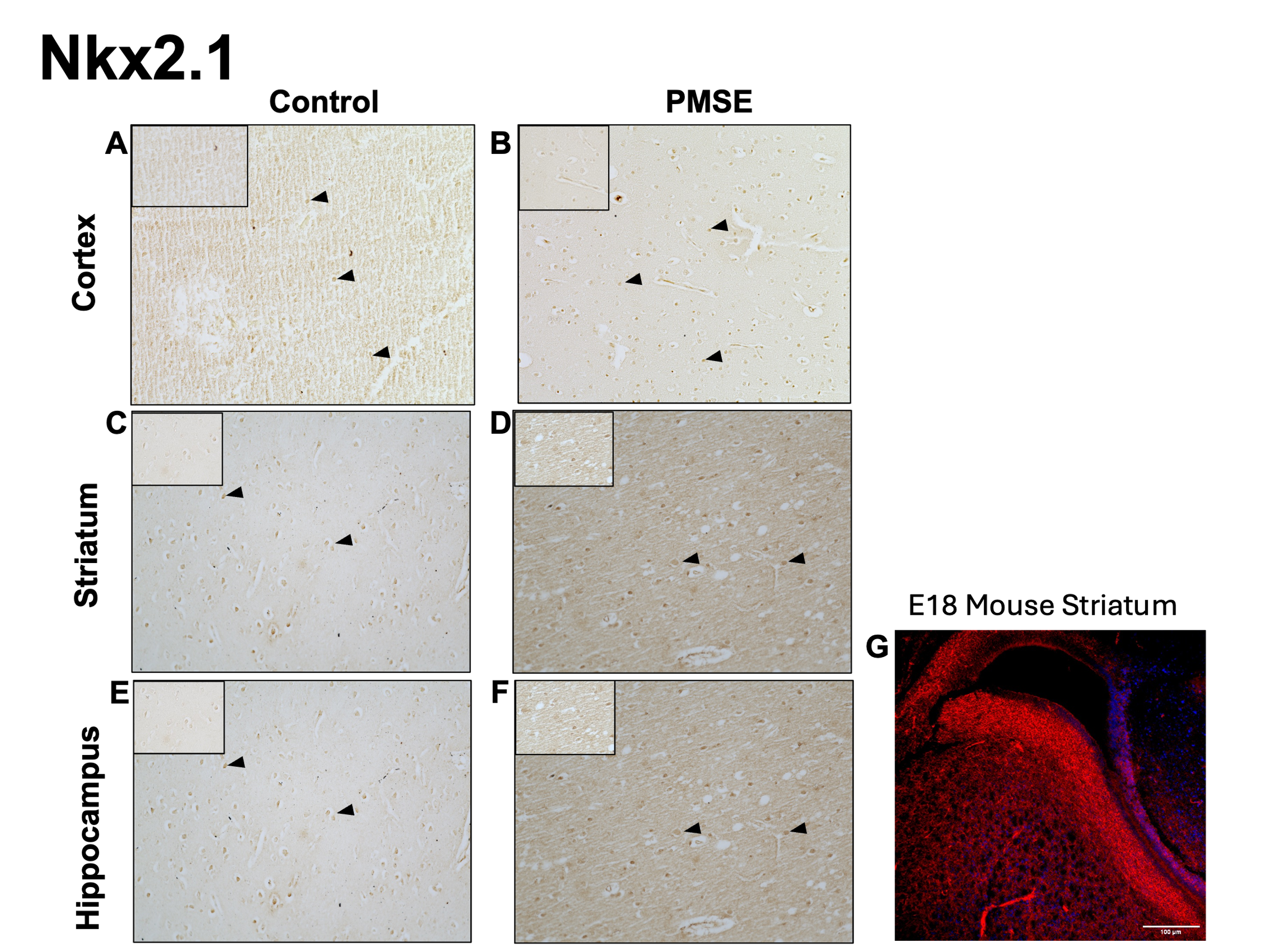
**Supplemental Figure 5:** No change in NKX2.1 labeling (**A-F**) in human control and PMSE tissue. Very little expression is observed across condition or brain region. Image is taken at 20x, inset is 40x with zoom factor of 1.6. This is supplemented by fluorescent MGE specific labeling of Nkx2.1 in embryonic day (E) 18 mouse tissue (**G-H**) using the same antibody to ensure that the lack of NKX2.1 expression in human tissue was not due to a poorly functioning antibody.

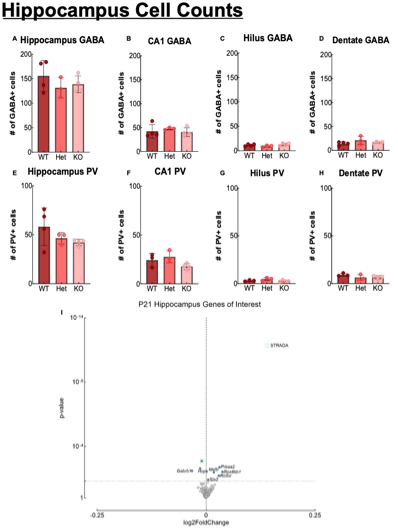
**Supplemental Figure 6**: Comprehensive hippocampus analysis of the expression of GABA+ (**A-D**) and PV+ (**E-H**) neurons including subregions (CA1, Hilus, DG) reveal no change across genotypes. RNA-seq in the hippocampus also reveal minimal changes in gene expression between WT and KO .

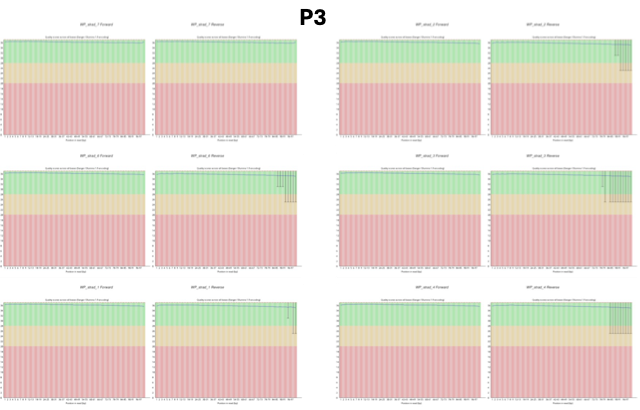

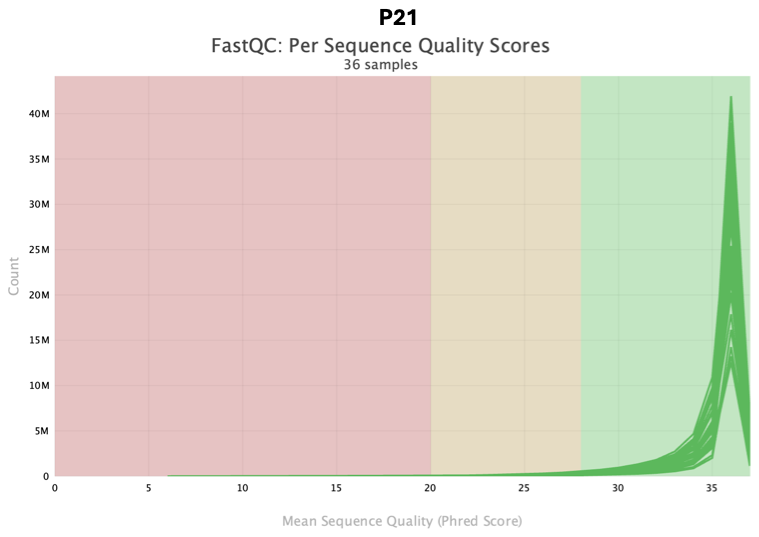

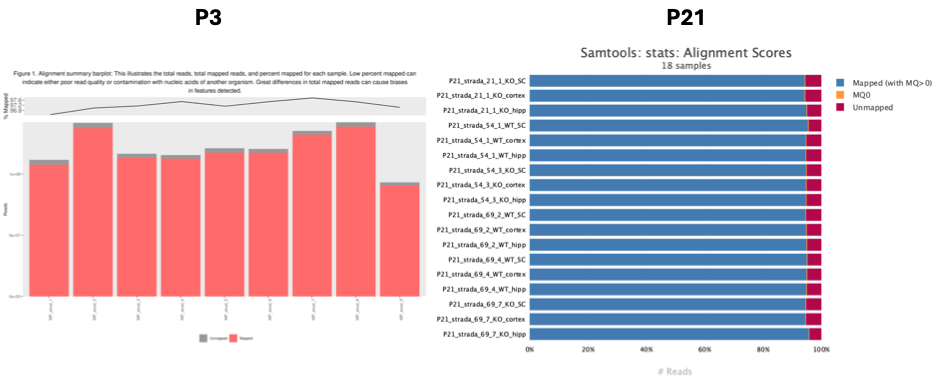

**Supplemental Figure 7:** Quality Control Report for P3 Cortex (**A,C**) separated by sample and P21 mice (**B,D**), with samples combined onto one graph. Each sample is a single animal. Generated by the Institute for Genome Sciences: The FastQC report presents graphical displays of the sequence quality. These graphs were created using the quality control software FastQC (Documentation: <https://www.bioinformatics.babraham.ac.uk/projects/fastqc/>). This report includes up to two samples per group. There are two graphs for each fastq file. The BoxWhisker plots illustrate the distribution of per base quality across all reads for the samples. The central red line is the median value, the yellow box represents the inter-quartile range (25-75%), the upper and lower whiskers represent the 10% and 90% points and the blue line represents the mean quality. The y-axis on the graph shows the quality scores. A higher score indicates a better the base call.

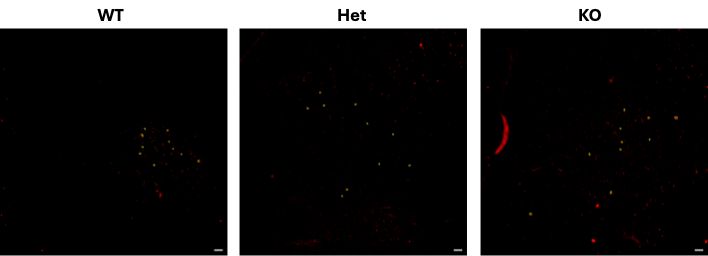

**Supplmental Figure 8**: Evidence of cytomegaly in GABA+ striatal neurons. Soma size traced and area measured in ImageJ of 10 representative cells in 3 sections per animal. Scale bars are 100µm.

**SUPPLEMENTAL TABLE LEGENDS**

**Supplemental Table 1**: Details of antibodies used for mouse and human immunolabeling including antibody, company, catalog number, concentration, and host species.

**Supplemental Table 2**: Semi-quantifications of human tissue using ImageJ automatic particle analysis

**Supplemental Table 3**: Mouse immunohistochemistry statistical analysis details

**Supplemental Table 4**: Master RNA-seq matrix of curated genes of interest list

**Supplemental Table 5**: Significant differentially expressed gene (DEG) list filtered by p-value <.05
